# 50 shades of greenbeard: Robust evolution of altruism based on similarity of complex phenotypes

**DOI:** 10.1101/2022.05.26.493612

**Authors:** Linnéa M Båvik, Rohan S Mehta, Daniel B Weissman

**Affiliations:** Department of Physics, Emory University

**Keywords:** altruism, cooperation, evolutionary dynamics, evolutionary game theory, mathematical biology

## Abstract

We study the evolution of altruistic behavior under a model where individuals choose to cooperate by comparing a set of continuous phenotype tags. Individuals play a donation game and only donate to other individuals that are sufficiently similar to themselves in a multidimensional phenotype space. We find the generic maintenance of robust altruism when phenotypes are multidimensional. Selection for altruism is driven by the coevolution of individual strategy and phenotype; altruism levels shape the distribution of individuals in phenotype space. Low donation rates induce a phenotype distribution that renders the population vulnerable to the invasion of altruists, whereas high donation rates prime a population for cheater invasion, resulting in cyclic dynamics that maintain substantial levels of altruism. Altruism is therefore robust to invasion by cheaters in the long term in this model. Furthermore, the shape of the phenotype distribution in high phenotype dimension—which is potentially more biologically relevant than low phenotype dimension—allows altruists to better resist the invasion by cheaters, and as a result the amount of donation increases with increasing phenotype dimension. We also generalize previous results in the regime of weak selection to two competing strategies in continuous phenotype space, and show that success under weak selection is crucial to success under strong selection in our model. Our results support the viability of a simple similarity-based mechanism for altruism in a well-mixed population.

## Introduction

An altruistic trait does not benefit the carrier of the trait but benefits other individuals with which the carrier interacts. Altruism can succeed under a wide variety of specific conditions, but the specter of a cheating individual who fails to reciprocate is constantly looming, and long-term maintenance of altruism requires some sort of mechanism to prevent cheaters from winning. This tension between altruism and cheating is present in many behavioral strategies in the natural world, including in alarm calls [She], the sharing of secondary metabolites [West et al., 2007], eusociality [Queller and Strassmann, 1998], slime mold life cycles [Strassmann and Queller, 2011], bacterial mat formation [Rainey and Rainey, 2003], and the major transitions of evolution [May].

Through decades of research, a general rule has emerged [Hamilton, 1964, Eshel and Cavalli-Sforza, 1982, Queller, 1985, Lehmann and Keller, 2006, Fletcher and Doebeli, 2008, Tarnita et al., 2009, Kay et al., 2020]: altruism can succeed if the interaction structure between individuals sufficiently disproportionately confers cooperative benefits upon cooperators. Potential mechanisms of this interaction structure include interacting with relatives [Hamilton, 1964, Cavalli-Sforza and Feldman, 1978], repeatedly interacting with the same individuals [Trivers, 1971], playing the game multiple times in a row [Axelrod, 1984], and interacting with individuals that are nearby in space [Nowak and May, 1992, Rousset, 2004]. These mechanisms are by no means mutually exclusive; many of them can be in operation at the same time, and they can interact with each other [Akçay and Van Cleve, 2012, Van Cleve and Akçay, 2014].

One class of interactions that can lead to altruistic success is interaction with phenotypically similar individuals, a specific form of which is called the “greenbeard” effect [Dawkins, 1976]. This effect requires a gene that codes both for a particular distinctive “tag” trait—such as a green beard—as well as the tendency to cooperate with others who have that trait. If such a gene were to exist, then the effects of the altruistic behavior would be completely distributed to altruists, and altruism can succeed. Greenbeard models generally have two limitations. First, it seems unlikely that there would exist a gene that encodes for both the tag trait and the behavioral trait, though some examples have been found in nature [e.g. Keller and Ross, 1998, Queller et al., 2003, Sinervo et al., 2006, Smukalla et al., 2008]. Secondly, and more importantly, greenbeards are highly susceptible to invasion by cheaters who have the tag trait but do not themselves cooperate [Gardner and West, 2010]; once these individuals are introduced to the population by mutation, it is unclear how the cooperative behavior would survive in the long term.

One solution to these problems with greenbeards is to decouple the tag and the behavior. A simple mechanism proposed by Riolo et al. [2001] allowed individuals to choose to cooperate based on how similar they were to each other with respect to a one-dimensional, continuous tag. This tag evolves separately from the altruistic strategy that determines the minimum similarity required to cooperate. Their model demonstrated the ability of such a system to maintain altruistic behavior over the long term by exhibiting a periodic co-invasion dynamic of more cooperative individuals (with larger minimum similarity thresholds) being invaded by less cooperative individuals (with smaller minimum similarity thresholds) which are then eventually re-invaded by more cooperative individuals based around a different set of tags. These cycles of altruistic behavior have been termed “tides of tolerance” [Sigmund and Nowak, 2001].

Controversy around the particulars of Riolo et al. [2001]’s model [Roberts and Sherratt, 2002] led to the development of a minimal two-tag, two-strategy model by Traulsen and Schuster [2003]. This and subsequent models [Traulsen and Nowak, 2007, Traulsen, 2008] have confirmed the existence of this “tides of tolerance” behavior while providing restricted conditions for when to expect it—including an upward mutational bias towards altruism or under a narrow range of mutation rate values.

Organisms simultaneously observe many traits in conspecifics, and use those observations to inform their interactions. For example, kin recognition in *Hymenoptera* involves comparisons of complex cuticular hydrocarbon profiles [Sturgis and Gordon, 2012]. Other examples with multidimensional phenotype matching include, territoriality in the bacterium *Proteus mirabilis* [Wall, 2016], and kin recognition in plants [Anten and Chen, 2021], birds [Krause et al., 2012], and gut microbiota in *Drosophila* [Lizé et al., 2014]. These examples suggest that interaction across multiple phenotype dimensions is a widespread phenomenon across the tree of life and should be considered in models involving phenotype-based behavioral interaction such as this.

Here, we analyze a model in the spirit of Riolo et al. [2001], Traulsen and Schuster [2003], Traulsen and Nowak [2007], Traulsen [2008] but taking into account the fact that individuals can recognize multi-dimensional continuous tags—or phenotypes. Using this model, we demonstrate long-term maintenance of altruistic behavior that is robust to invasion by cheaters over a wide range of parameter values. Our model is similar to that of Spector and Klein [2007], which found an increase in donation with increasing phenotype dimension; crucially, however, our model differs from theirs in that we assume a well-mixed population, isolating the effects of phenotypic space from those of physical space. The robustness of altruism in our model is fundamentally driven by the shape of the population phenotype distribution in multidimensional phenotype space. In particular, in high-dimensional phenotype space, it is extremely rare for individuals to be similar across their entire phenotypes unless they share recent common ancestry. This phenotype distribution makes it less likely for non-altruistic individuals to find more than a small amount of altruistic individuals to cheat off of and dominate. In addition, we extend the weak-selection results of Antal et al. [2009] and Kroumi and Lessard [2015] to two competing strategies with nonzero, continuous thresholds, and we demonstrate that altruistic success in this weak selection setting is crucial to altruistic success in our strong selection setting. Our results suggest that the high dimensionality of phenotype space is sufficient to select for biased altruism in the absence of additional mechanisms such as mutation bias, spatial structure, or reciprocation.

## Model

Our model consists of a single, well-mixed population of *N* individuals, each with two traits: a visible phenotype and an invisible strategy, illustrated in Figure 1. We model the visible phenotypes as quantitative traits. An individual *i*’s phenotype, denoted Φ_*i*_ ∈ R^*D*^, is a point in a *D*-dimensional continuous phenotype space that can be observed by other individuals in the population. Individuals play a donation game with other individuals. An individual’s strategy, called its “threshold” *T*_*i*_, determines whether or not it will donate to another individual. An individual *i* will donate if the potential recipient *j* is sufficiently similar to them, i.e. the distance between the two individuals in phenotype space is less than *T*_*i*_: ||Φ_*i*_ − Φ_*j*_|| ≤ *T*_*i*_.

**Figure 1:**
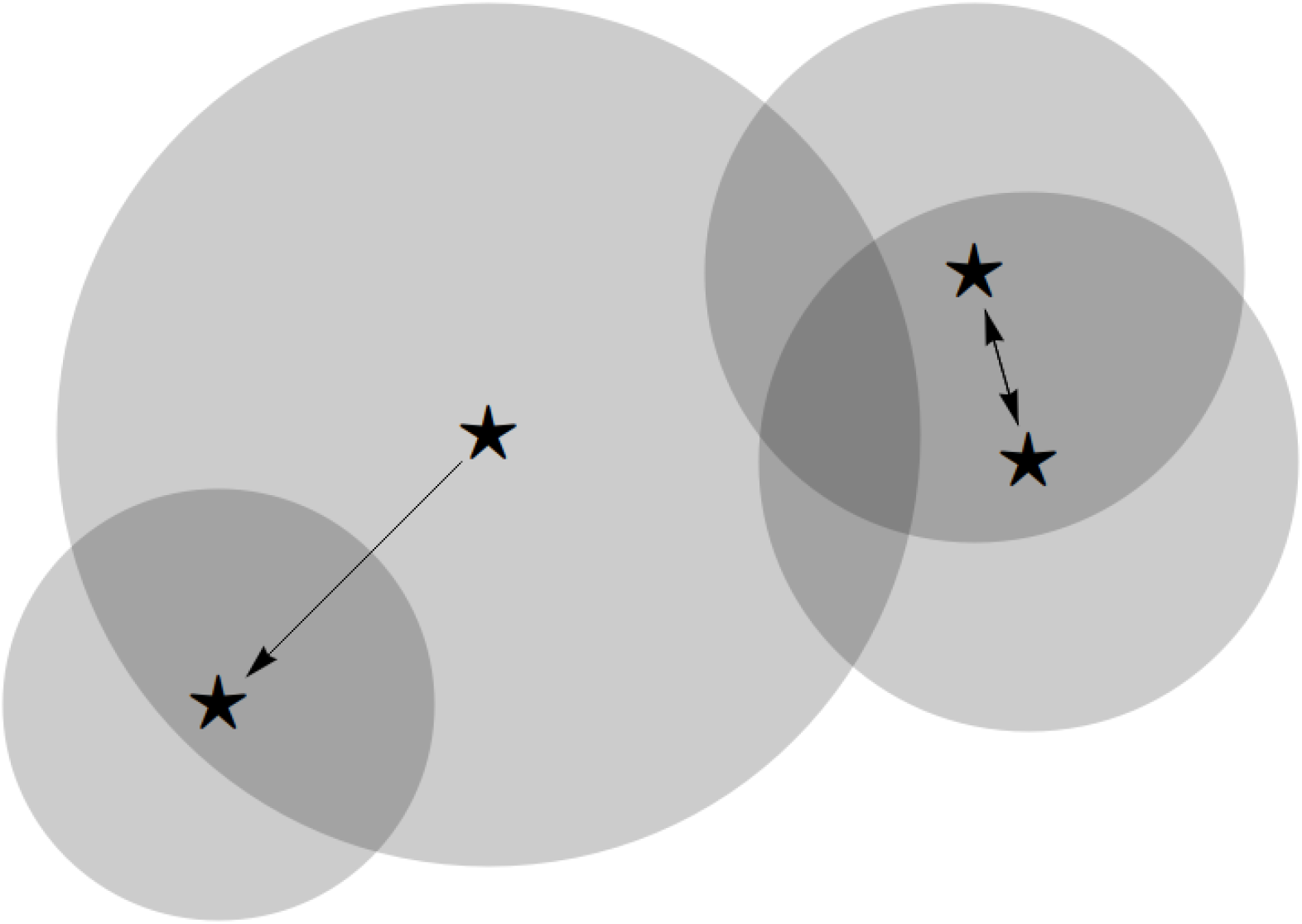
Schematic of the model for biased altruism. Individuals (stars) are shown as points in an abstract two-dimensional phenotype space. Each individual also has a threshold for altruism represented by a gray circle. They donate to all individuals within their circle, signified by arrows pointing from donors to recipients.

We assume uniparental reproduction, with offspring inheriting their parent’s phenotype and strategy, plus slight deviations from additive mutations. This could describe genetic inheritance in an asexually reproducing population, or cultural inheritance from one parent or other individual in the parental generation. Phenotype mutations occur with every reproduction and are drawn from a standard multivariate normal distribution *N* (0, **1**) where **1** is the *D*-dimensional identity matrix. In other words, we measure phenotypes on the natural scale set by mutation. Strategy mutations occur with probability *ν* ≪ 1 per reproduction and are drawn from a truncated normal distribution *TN* (0, *σ*_*S*_, −*T*_*i*_, ∞). We use a truncated distribution to avoid unbiological negative thresholds. The truncation creates a small upward mutational bias on the order of *σ*_*S*_ when the threshold *T*_*i*_ approaches 0, so we interpret thresholds on the order of *σ*_*S*_ to be noncooperative. In this model, the probability of strategy identity between parent and offspring is high but the probability of phenotypic identity is low—the opposite of Riolo et al. [2001].

In every generation, each individual observes ten potential donation recipients chosen at random from the rest of the population. If the distance between the individual and a potential recipient in less than the individual’s similarity threshold, the individual increases the recipient’s payoff by the donation value *b*. If the potential recipient is outside of the individual’s similarity threshold, the individual declines to donate, instead keeping their resource for themselves, thereby incrementing their own payoff value by *c*, where *c < b*. The fitness of each individual is proportional to the total payoff acquired by these donation events. Note that the payoff differentials incurred through this donation game are identical to the cost-benefit form of the payoff matrix of the prisoner’s dilemma, with the payoff values chosen to avoid negative numbers in the fitness weighting. For details about the mutation and selection processes in this model as well as the simulation procedure, see Appendix A.

## Results

### High phenotypic complexity leads to the evolution of frequent altruism

We find that for low phenotypic dimension *D* ≲ 10, altruism rarely evolves and donations are rare. But for high phenotypic dimension *D* ≳ 10, altruism consistently evolves and donations are maintained at a substantial rate of about 10-25%, so that in each generation, an individual donates to about one to two of the ten others they interact with (Figure 2). For each particular phenotype dimension *D*, the threshold evolves to a narrow evolutionarily stable nonzero range of values (Figure 3). For lower-dimensional phenotypes, there are regular oscillations within this range, but for higher-dimensional phenotypes these disappear. This pattern is robust to changes in population size, payoff values, mutation rate, initial conditions and phenotype dimension, although the specific value of the evolutionarily stable threshold is a function of phenotype dimension and population size (Figure 3 and Appendix B).

**Figure 2:**
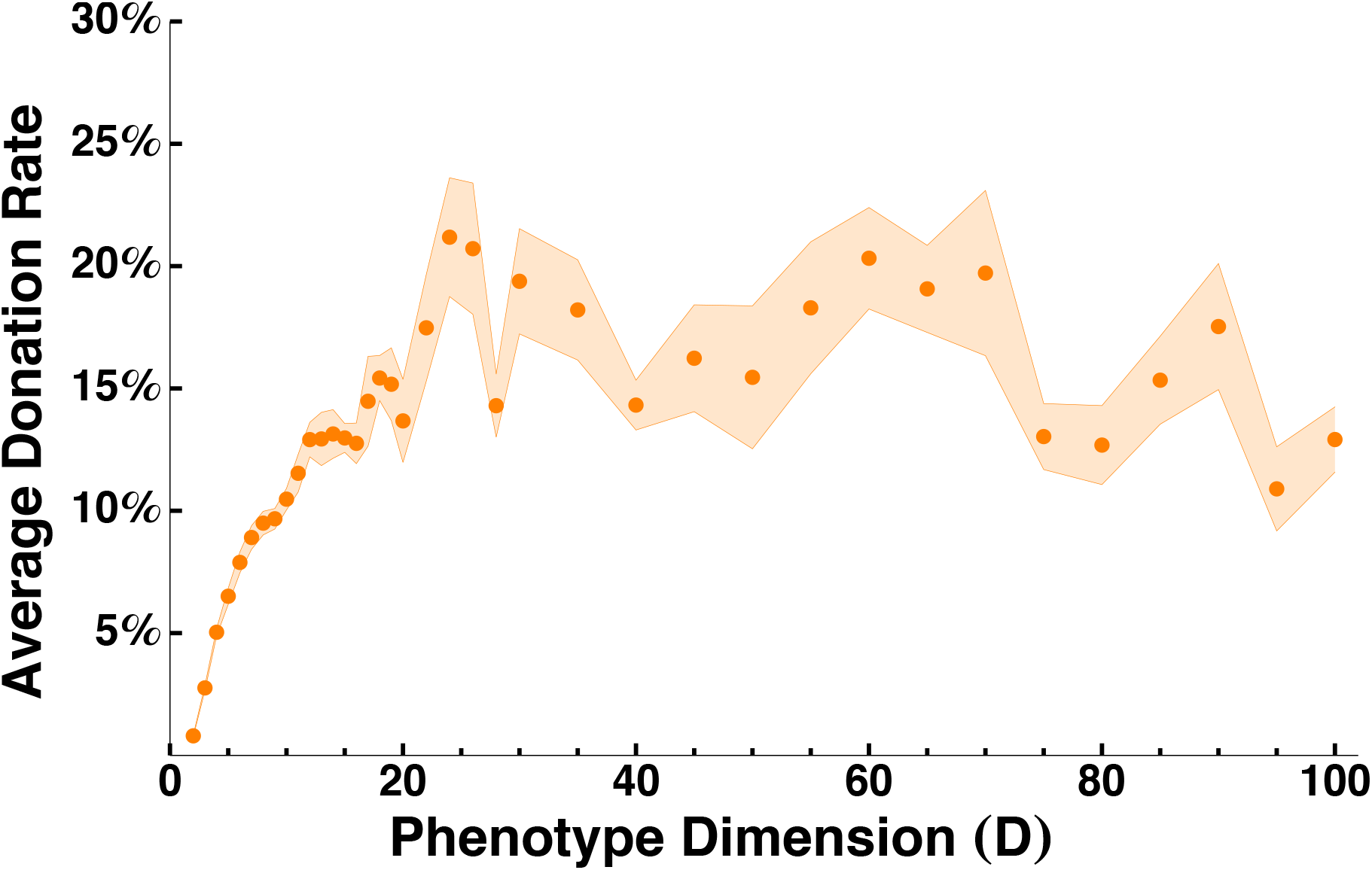
High-dimensional phenotypes lead to the evolution of altruism. Donation rate is the fraction of possible donations that occur. For each simulated dimension, two different initial conditions were run for five iterations each, for a total of ten simulations. Points show the average across these ten simulations, with ribbon indicating the standard error. For each simulation, we took the average donation rate of the last 100,000 generations, sampled every 100 generations. Parameters used in these simulations are in Table 1.

**Figure 3:**
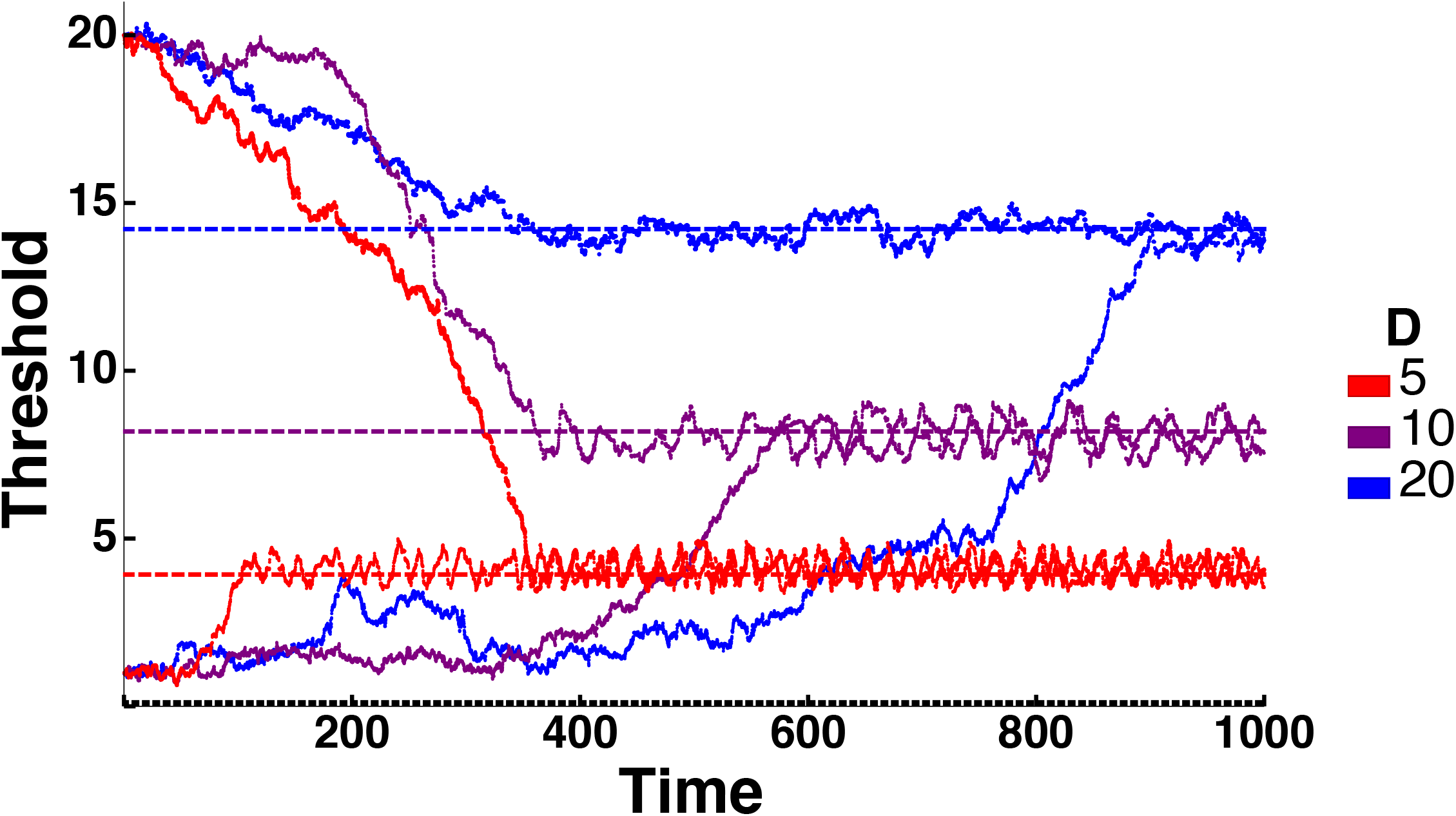
Altruism thresholds evolve to high values that increase with phenotype dimension, indicating the maintenance of altruism in the long term. Each line shows a single simulation, with the two lines for each color starting from two different initial conditions. Dashed lines show the cohesion thresholds as identified in Figure 4. Dotted line shows *σ*_*S*_, the characteristic threshold scale in non-altruistic populations. Parameters used in these simulations are in Table 1. Time is in units of *N* = 1000 generations of the simulation.

### Altruism shapes the distribution of phenotypes

By simulating populations with no strategy mutation, where each individual has the same strategy, we found that there is also a characteristic threshold in these single-strategy evolutionary dynamics (Figure 4). We refer to it as the “cohesion threshold”, because it causes the population to cohere in a small region of phenotype space. Figure 4B illustrates that for smaller thresholds, donation is very rare, and so there is little variation in individual fitness. Thus, the phenotypic variation is nearly neutral, with a variance predicted by coalescent theory (see the dashed horizontal line in Figure 4A and Appendix D).

**Figure 4:**
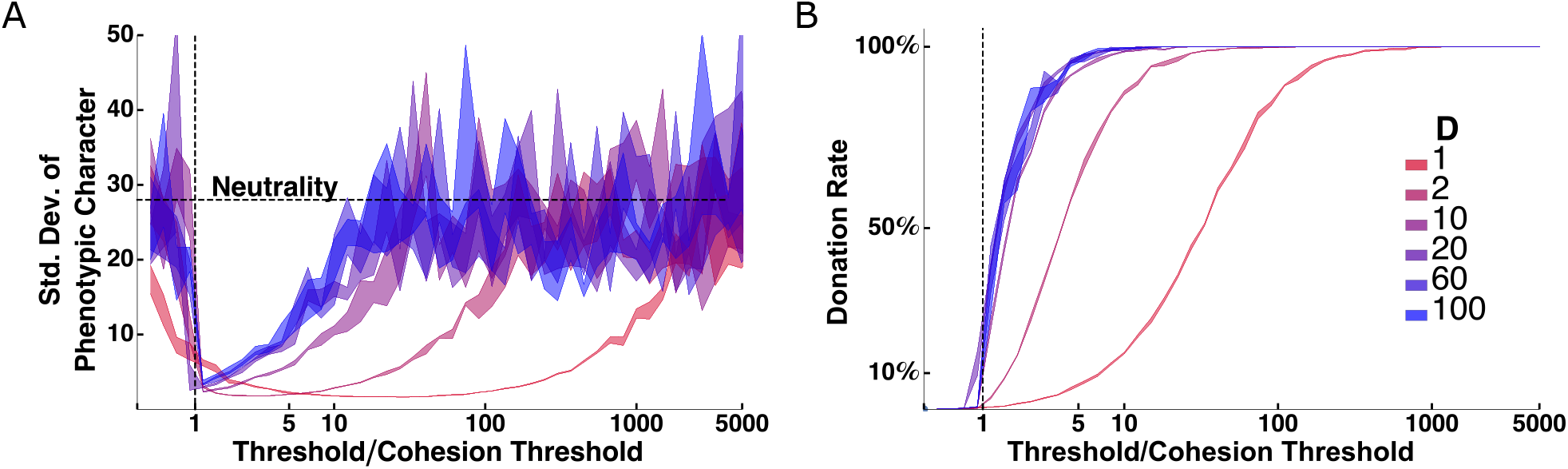
There is a critical “cohesion threshold” at which altruism greatly reduces equilibrium phenotypic diversity. Curves show the results of simulations in which all individuals have a single fixed threshold, shown on the horizontal axis (scaled by the cohesion threshold). (A) The standard deviation of each phenotypic dimension collapses to a small value at the cohesion threshold, and increases gradually as threshold increases beyond the cohesion threshold. For thresholds below and far above the cohesion threshold, the phenotype distribution approaches the distribution expected under neutrality (dashed horizontal line, Appendix D). (B) The cohesion threshold is the threshold at which donation starts to become frequent. Ribbons indicate the standard deviation of the measured quantity (phenotype standard deviation or donation rate) over all iterations. Parameters used in these simulations are in Table 1.

When thresholds are far above the cohesion threshold, donation is universal and phenotypic variation is again neutral. But at and somewhat above the cohesion threshold, there is an intermediate amount of donation, creating a large variance in fitness sufficient to counteract mutation pressure and greatly reduce phenotypic variation. While the cohesion threshold appears for all dimensions, donation is much more prevalent at the cohesion threshold for high-dimensional phenotypes than for low-dimensional ones (Figure 4B).

For high-dimensional phenotypes, the cohesion threshold increases with increasing population size (Appendix C). This effect appears to be less pronounced as the population size increases, suggesting that there is an infinite-population-size asymptotic value to the cohesion threshold.

### Coevolution of strategies and phenotypes maintains altruism

Remarkably, it appears the two independently identified characteristic thresholds, the evolutionarily stable threshold and the cohesion threshold, are the same (Figures 3 and 5). In other words, populations evolve to just the threshold at which their phenotype diversity is narrowest. For high-dimensional phenotypes (*D >* 20), there is an additional convergence, as the width of the population diversity in each phenotypic dimension also approaches the same value as the two thresholds (Figure 5); this is what gives rise to the substantial frequency of donations seen in Figure 2.

**Figure 5:**
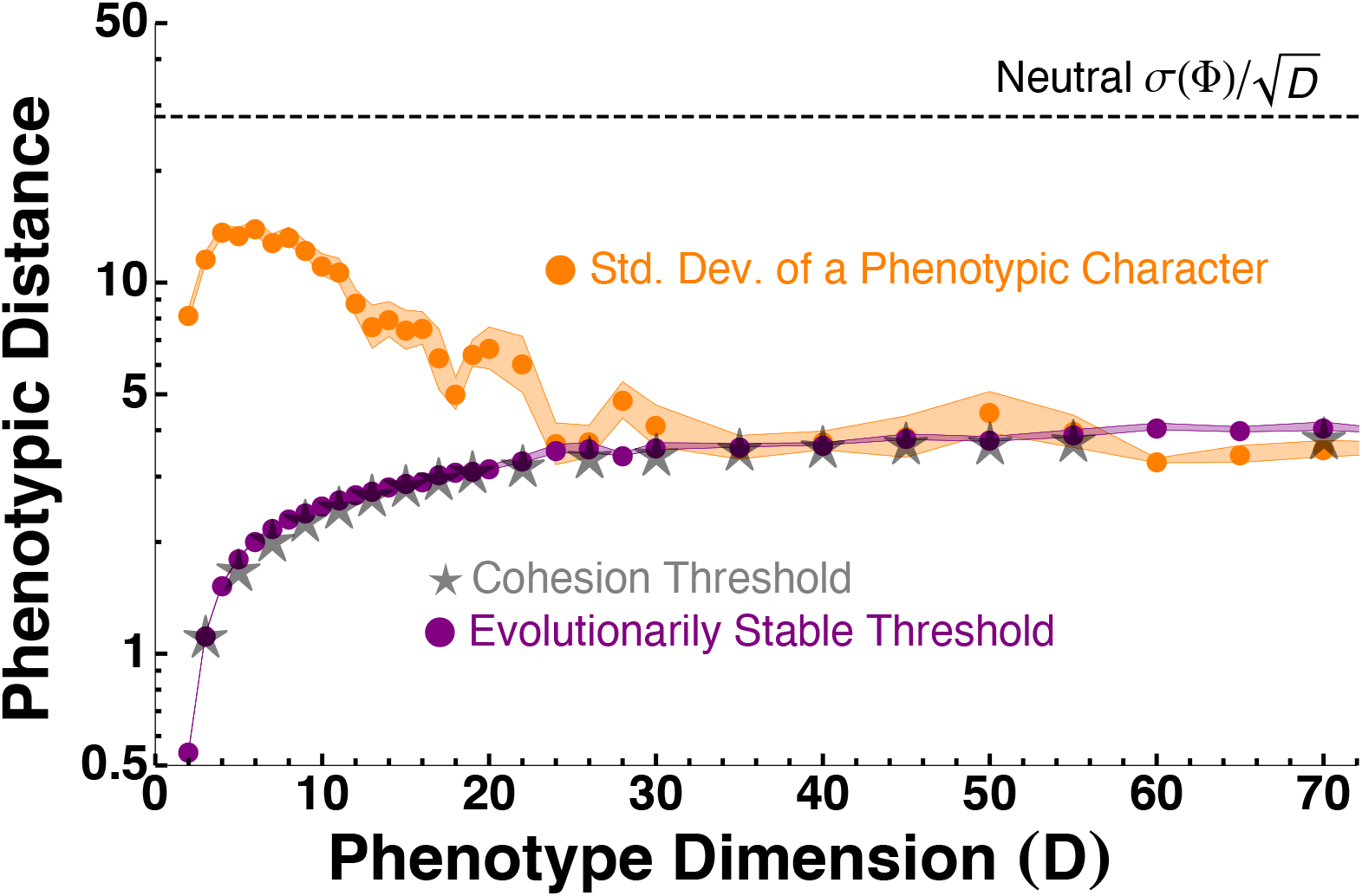
The cohesion threshold (gray) is the evolutionarily stable threshold (purple). For high-dimensional phenotypes, the standard deviation of phenotypic diversity (orange) converges to this value as well. All values are scaled by √*D* to obtain the value along each phenotypic dimension. The evolutionarily stable threshold and phenotype standard deviation are found using simulations of evolving thresholds, while the cohesion threshold is found using simulations with fixed thresholds. Parameters used in these simulations are in Table 1.

To understand why the cohesion threshold is evolutionarily stable, first consider a population dominated by individuals with thresholds smaller than the cohesion threshold. Then donation is rare (Figure 4B) and only slightly perturbs the distribution of phenotypes, which is close to neutral (Figure 4A). This is the key assumption underlying the standard weak selection models in evolutionary game theory (see Appendix D). Under weak selection, the larger threshold will succeed if the benefit/cost ratio of donation is greater than some critical value. For our simulations, we have chosen *b/c* = 10, as in Riolo et al. [2001], which is high enough to produce this selective pressure under most circumstances, causing the population to evolve up towards the cohesion threshold. We conduct a mathematical analysis of this regime in Appendix D, generalizing the results of Antal et al. [2009] and Kroumi and Lessard [2015] to continuous strategies.

Now consider a population where most individuals have thresholds substantially above the cohesion threshold, so that donation is very high—most individuals donate to most individuals (Figure 4). In this situation, individuals with smaller threshold values can invade, since most other individuals will still donate to them, driving the threshold down towards the cohesion threshold. This process can only stabilize once there is frequent discrimination, i.e., when thresholds are low enough that many interactions do not result in donation.

### High-dimensional phenotypes protect altruists from invasion by cheaters

Figures 2 and 5 show a pattern in which low-dimensional phenotypes produce modest donation rates, moderate phenotypic diversity, and cycling about the cohesion threshold, while high-dimensional phenotypes produce high donation rates, low phenotypic diversity, and little cycling. To understand the origins of this pattern, we simulated populations in which individuals mutated between just two discrete thresholds, one below the cohesion threshold (“strict”) and one above it (“generous”). These simulations showed that the different features are the result of a single process: low-dimensional phenotypes allow strict individuals to invade generous populations, because they are tightly clustered in phenotype space (Figure 6, time 1). This then causes the population to decohere and spread out in phenotype space (time 2), producing the elevated average phenotypic diversity seen in 5. The phenotypic diversity allows generous individuals to re-invade from a sparse region of phenotype space where they avoid strict cheaters (time 3). But these generous individuals re-collapse the phenotype distribution, resetting the cycle (time 4).

**Figure 6:**
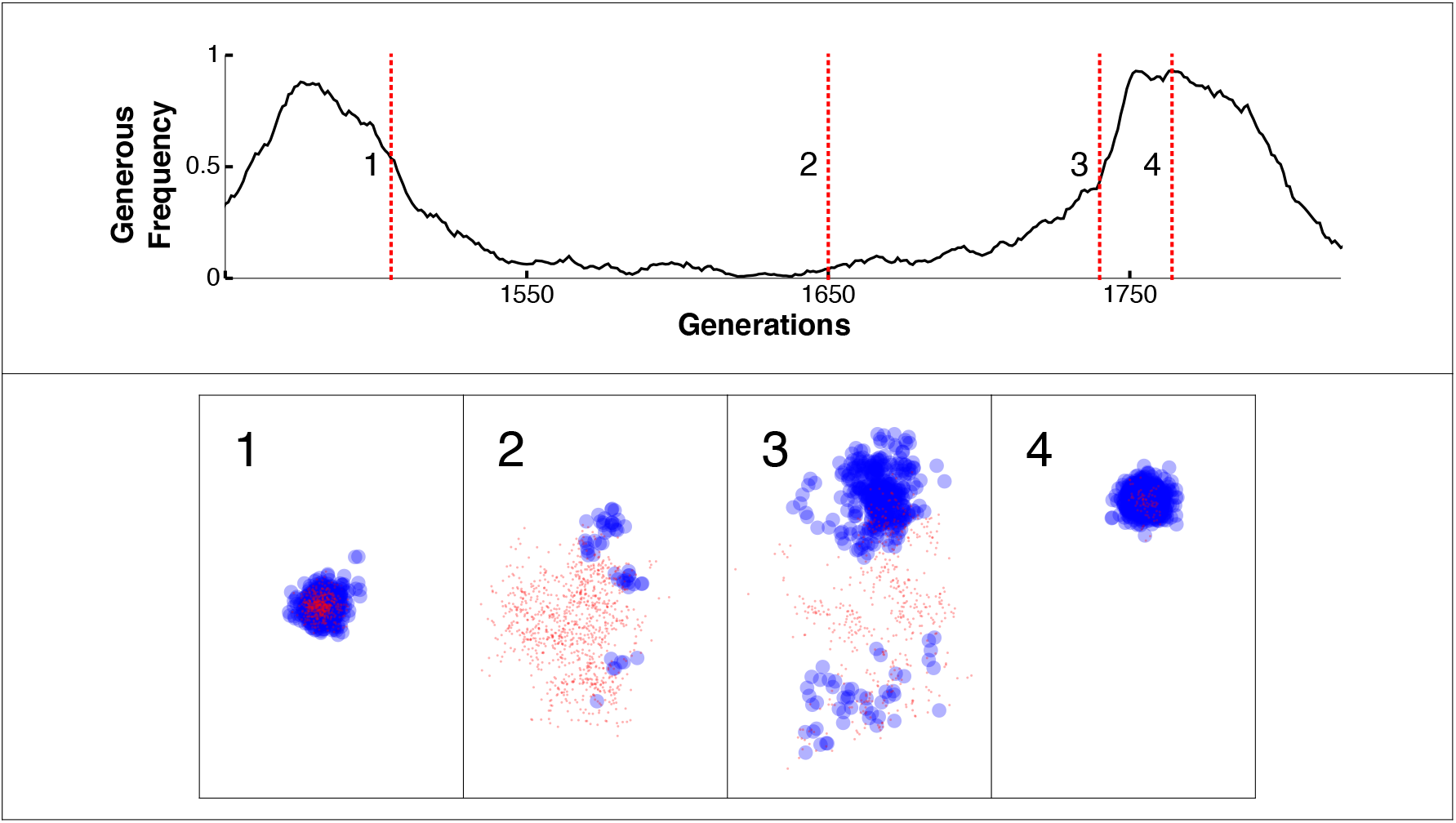
Low-dimensional phenotypes allow non-altruists to invade. Plots show results of simulations of a two-dimensional phenotype over the course of a cycle similar to those seen for *D* = 5 and 10 in Figure 3. For clarity, individuals are restricted to only two possible thresholds, one strict and one generous—corresponding to thresholds below and above the cohesion threshold, respectively. Red circles are “strict” individuals, blue circles are “generous” individuals, with the radii of the circles equal to their thresholds. Time point 1: strict individuals have started to invade the cohesive cluster of generous individuals. Time point 2: the strict individuals have successfully invaded, causing the population to disperse in phenotype space. Time point 3: generous individuals from the sparse edge of the phenotype distribution re-invade. Time point 4: the population re-coheres in phenotype space and is again vulnerable to invasion by strict individuals. Parameters used in this simulation are in Table 1.

High-dimensional phenotypes, in contrast, do not allow strict individuals to easily invade. This may seem surprising, given that Figure 5 shows that they have limited phenotypic variation, but it is simply a consequence of geometry: high dimensions leads to a high surface-to-volume ratio for the phenotype distribution, with essentially all individuals living on the sparse “edge” of the distribution. While strict individuals can invade locally in phenotype space they cannot reliably spread to parasitize the rest of the population, and so quickly become isolated and go extinct. The population thus stays cohesive with frequent donations.

## Discussion

Our model shows that high-dimensional phenotypes allow the evolution of biased altruism by mitigating invasion by cheaters: high-dimensional space is large enough to ensure that phenotypic similarity is an honest signal of close relatedness. Previous work on the evolution of altruism by phenotypic similarity established relatively stringent conditions on mutation rates, the structure of phenotype space, or the structure of the population in physical space required for altruism to succeed. Our model demonstrates that this success can exist quite generally, independent of the particulars of the mutation rates. In addition, our model shows that with high-dimensional phenotypes, altruism does not need any particular population structure or individual memory or even reciprocal pairwise interactions between individuals to succeed.

It has often been difficult to determine the utility of weak selection models outside of the weak selection limit. Our model demonstrates that there are occasions where such analysis can provide insight outside of that limit. It is therefore worth analyzing models that have been only formulated in the weak selection limit outside of this limit to see how they can inform our understanding of scenarios with stronger selection.

The phenotypic space within which our simulated individuals live is distinct from the physical space in which they would live in two primary ways. First, physical space is limited to no more than three dimensions, whereas phenotype space often has many more dimensions. The high number of dimensions is crucial to the evolution of altruism in our model, so we predict that phenotype-based altruism can be maintained even when space-based altruism cannot. Second, in the functionally neutral phenotype space of the type we consider, there is no local density regulation—individuals do not compete more for resources with phenotypically similar individuals than they do with phenotypically distant individuals. This distinction allows for the differences in local density that drive our results to be unaffected by considerations of local competition that influence models of altruism based on proximity in physical space (see, e.g., Taylor [1992]’s famous cancellation result).

In actual biological populations, there are many possible phenotypic traits that organisms can observe or detect in making behavioral decisions towards other organisms. In many cases, the traits involved in processes like kin recognition involve multiple loci and are highly chemically diverse. We argue that “realistically-sized” phenotype spaces of many tens to hundreds of dimensions are a more appropriate setting for the understanding of the evolution of behavior based on phenotypic similarity. In particular, the fact that high-dimensional phenotypes imply that everybody is atypical in some way is a general phenomenon and may be relevant to many more aspects of behavioral evolution is beyond the evolution of altruism. Thus, altruism based on phenotypic similarity is much less constrained by the properties of the space in which the strategy exists than altruism based on physical proximity.

A potential avenue for further study would be to also allow the phenotype dimension to evolve to see if altruism can drive or limit the evolution of phenotypic complexity and the complexity of phenotypic sensing. It would also be interesting to consider strategies that do not use a fixed threshold but instead learn their threshold based on the current phenotype distribution. Finally, it is important to consider the potential effects of different forms of phenotypic transmission. Most obviously, one could consider a sexually reproducing population, possibly with assortative mating or inbreeding depression. But it is also important to consider the possibility that the phenotype and strategy may be cultural traits with potential horizontal transmission.

## A Methods

### Selection

At every time step, every individual chooses whether or not to donate with ten other individuals chosen uniformly at random from the population, and the total payoff resulting from each of these games weights the probability of being chosen in the next generation under the Wright-Fisher process by the following equation:

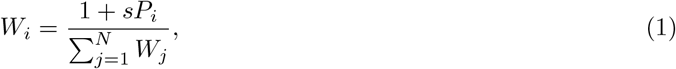

where *W*_*i*_ is the weight of individual *i* in the multinomial draw for the next generation in the Wright-Fisher process, *P*_*i*_ is the total payoff of individual *i* after playing the game with every other individual, and *s* is a parameter that describes the strength of selection. For large *s*, selection is strong, and for *s* → 0 we obtain the weak selection limit (see Appendix D). For the this analysis, we set *s* = 1 for simplicity so that the difference in weight is equal to the difference in payoff.

### Simulation

For any particular set of parameters, we simulate a population of *N* = 1000 individuals where each time step is a generation in the Wright-Fisher process. At each time step, every individual chooses whether or not to donate to ten other individuals chosen uniformly at random from the population. The simulation code is available at https://github.com/weissmanlab/Robust evolution of altruism based on similarity of complex phenotypes. Parallelization of this code used GNU Parallel [Tange, 2011].

Table 1: Model parameters. The number of generations varied for Figures 2 and 5 because higher dimensions required longer time to equilibrium. *CT* = cohesion threshold

**Table 1.** lists the parameters we use in our simulations for each figure.

| Symbol | Description | Range | Figures 2 and 3 | Figure 4 | Figure 6 | Figure 5 | Figure 7 | Figure 8 |
| --- | --- | --- | --- | --- | --- | --- | --- | --- |
| $N$ | Number of individuals in population | $[1, \infty)$ | 1000 | 1000 | 1000 | 1000 | 1000 | 1000 |
| | Number of generations for simulation | $[1, \infty)$ | $3 \times 10^5 - 3 \times 10^6$ | 3000 | 4000 | $3 \times 10^5 - 3 \times 10^6$ | $3 \times 10^5$ | 1100 |
| $b$ | Benefit of altruism | $[0, \infty)$ | 10 | 10 | 10 | 10 | 10 | 10 |
| $c$ | Cost of altruism | $[0, \infty)$ | 1 | 1 | 1 | 1 | 1 | 1 |
| $T$ | Threshold for altruism | $[0, \infty)$ | N/A | Fixed | N/A | N/A | N/A | N/A |
| $T_G$ | Generous (bigger of two) thresholds | $[0, \infty)$ | N/A | N/A | 1.5 | N/A | N/A | N/A |
| $T_S$ | Strict (smaller of two) thresholds | $[0, \infty)$ | N/A | N/A | 0.25 | N/A | N/A | N/A |
| $s$ | Strength of selection | $[0, \infty)$ | 1 | 1 | 1 | 1 | 1 | 1 |
| $PM$ | Standard deviation of phenotype mutation distribution | $[0, \infty)$ | 1 | 1 | 1 | 1 | 1 | 1 |
| $\nu$ | Probability of strategy mutation | $[0, 1]$ | 0.001 | 0 | 0.0001 | 0.001 | 0.1, 0.01, 0.001 | 0 |
| $\sigma_S$ | Standard deviation of strategy mutation kernel | $[0, \infty)$ | 0.1 | N/A | N/A | 0.1 | 0.001 | N/A |
| $D$ | Phenotype dimension | $\mathbb{Z}^+$ | 1-100 | 1-100 | 2 | 1-70 | 2, 10, 50 | 1-125 |
| Initial $T$ | Value of $T$ at time $t = 0$ | $[0, \infty)$ | $0.75 \times CT, 1.25 \times CT$ | Varies | N/A | $0.75 \times CT, 1.25 \times CT$ | $0.5 \times CT, 1.5 \times CT$ | Varies |

### Determination of cohesion threshold

Intuitively, the cohesion threshold is the threshold at which the equilibrium phenotypic diversity drops precipitously. We operationalized this definition for our specific parameter values as the lowest threshold at which the standard deviation per phenotypic dimension fell below 5 (i.e., 5 times the mutational effect size). To find this threshold, we used a lattice of simulated threshold values centered on the predicted cohesion threshold for each value of the dimension *D*.

## B Robustness to parameter variation

Figure 7 displays the temporal dynamics of strategy evolution for various different parameter values (see Table 1 for the specific values.) In each case, for two different initial conditions starting above and below the cohesion threshold, the strategy evolves towards the cohesion threshold. Increasing the strategy mutation rate *ν* speeds up the approach to the cohesion threshold, and increasing phenotype dimension *D* slows down the approach to the cohesion threshold.

**Figure 7:**
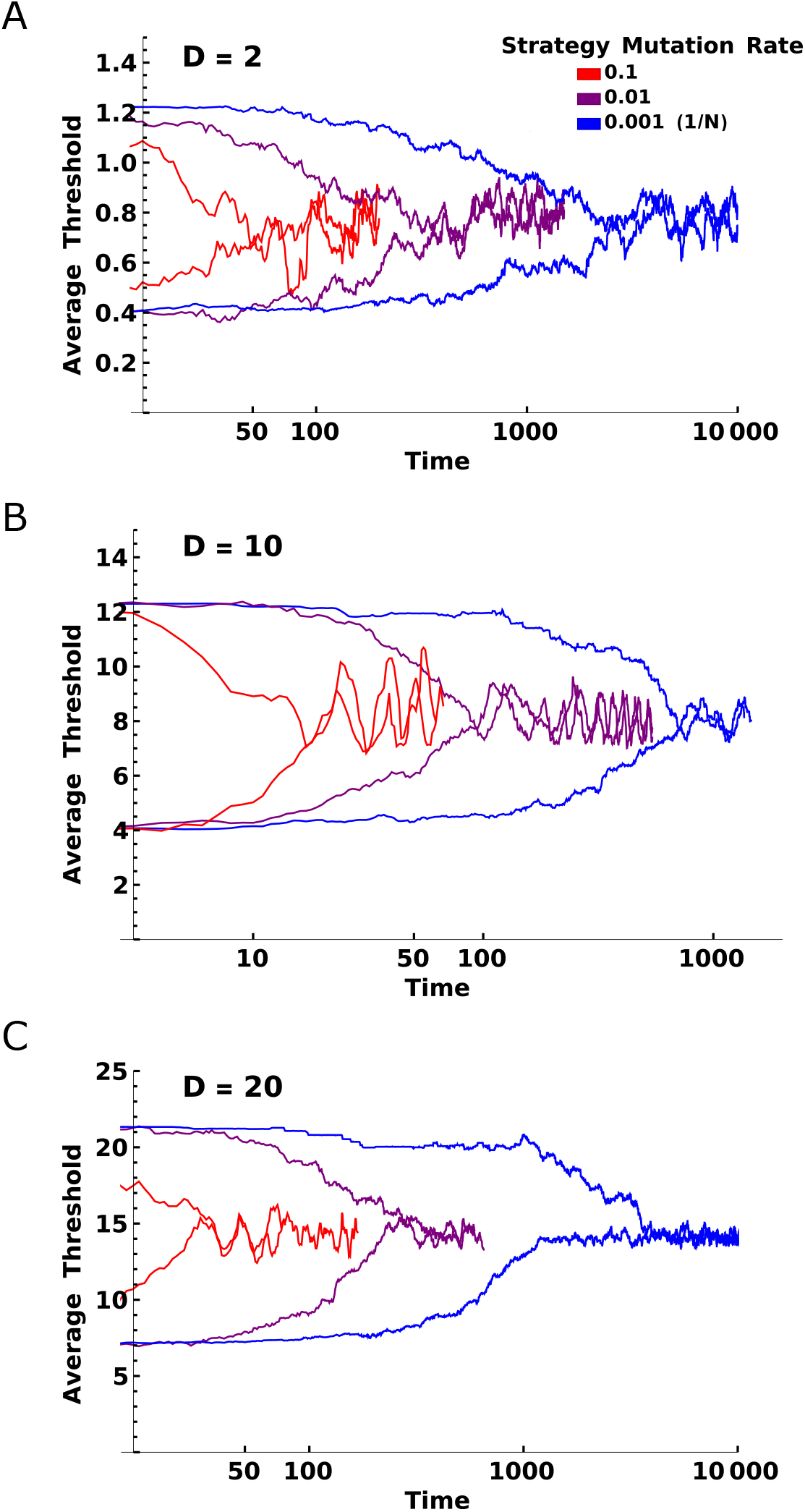
Evolution to the cohesion threshold is robust to variation in phenotype dimension and strategy mutation rate (*ν*). Each panel shows, for two different initial conditions, the threshold evolving to the cohesion threshold, terminating when the two initial condition trajectories meet. Parameters used for these simulations are summarized in Table 1, with the exception of *D* = 2, where *σ*_*S*_ = 0.01 to make sure that the cohesion threshold was significantly above the mutation parameter.

## C Effect of population size on cohesion threshold

Figure 8 displays the value of the cohesion threshold for different values of the population size *N*. In each case, cohesion threshold increases with population size, and this effect is most pronounced for very high phenotype complexity. The pattern suggests that populations stay close to a deterministic (i.e., infinite *N*) limit up to a certain limiting phenotype dimensionality, with that limiting dimensionality increasing with population size.

**Figure 8:**
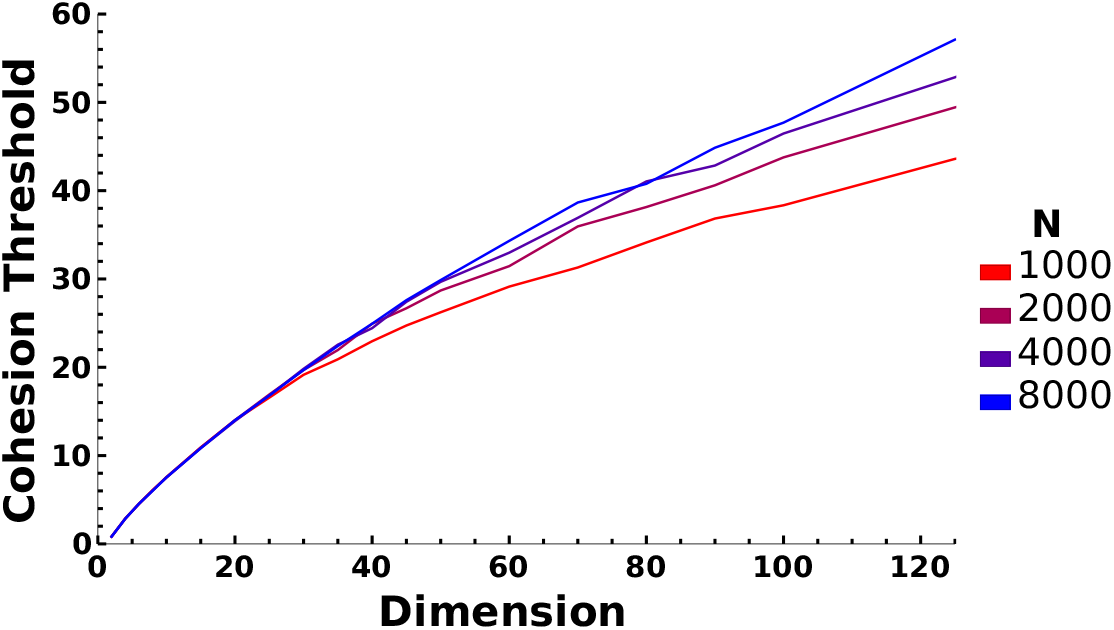
Cohesion threshold increases with increasing population size, with the effect only notable for large phenotype dimension. Simulation parameters are in Table 1.

## D Analysis for weak selection

The phenotype distribution when all thresholds are below the cohesion threshold resembles a neutral phenotype distribution (Figure 4). The assumption that the phenotype distribution is unaffected by selection is precisely the condition needed for a mathematically tractable weak selection analysis. To rigorously obtain the expectation of the phenotype distribution under neutral assumptions, and to obtain a prediction of the success of one strategy compared to another under the model of weak selection, we proceed with a full analysis of our model under the assumption of weak selection. Formally, we let *s* → 0 in (1) in a population of finite size with two strategies: one generous and one strict. A generous individual has a threshold *T*_*G*_, and a strict individual has a threshold *T*_*S*_, with *T*_*G*_ *> T*_*S*_. Individuals can mutate between the two strategies with probability *u* and undergo phenotype mutation as described in the Model section. In this case, we allow individuals to choose whether or not to donate to the entire rest of the population as opposed to just ten other individuals.

Much of the analysis in this section follows Antal et al. [2009] and Kroumi and Lessard [2015], so we will merely restate these previous results until they differ from ours. Following Antal et al. [2009], Kroumi and Lessard [2015], we consider the condition for success of the generous strategy under weak selection to be that the expected frequency of generous individuals in the stationary state of the population is greater than 1*/*2. Eq. 10 of Kroumi and Lessard [2015] yields this expected frequency

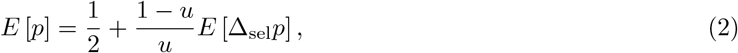

where *u* is probability of strategy mutation and *E* [Δ_sel_*p*] is the expected change in the frequency of generous individuals due to selection. To obtain the frequency of generous individuals in the stationary state of the system, we must compute *E* [Δ_sel_*p*].

Call *a*_*ϕ*_ the number of generous individuals at phenotype *ϕ*, and *b*_*ϕ*_ the number of strict individuals at phenotype *ϕ*. Let us also define

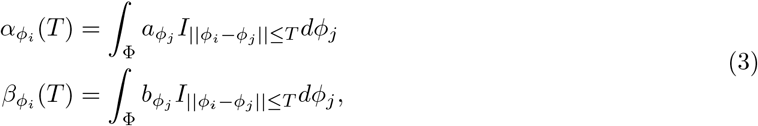

to be the number of generous and strict individuals, respectively, within a distance *T* of phenotype *ϕ*_*i*_ (with *I* being the indicator function for this condition). Call P the set of phenotypes in the population. We also define *n*_*ϕ*_ = *a*_*ϕ*_ + *b*_*ϕ*_ and *N*_*ϕ*_(*T*) = *α*_*ϕ*_(*T*) + *β*_*ϕ*_(*T*). The payoffs of a generous individual and a strict individual at phenotype *ϕ*_*i*_ are therefore

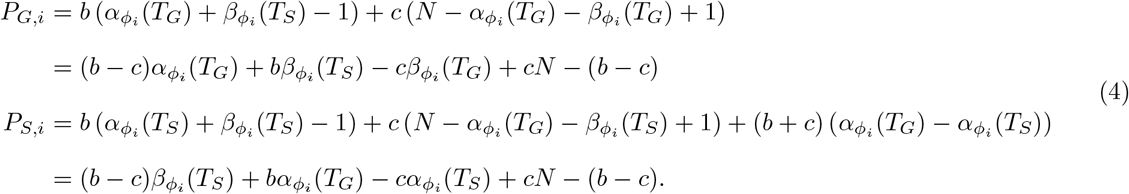

where we have corrections for disallowing self-interaction in the form of subtracted 1s.

The corresponding selection-adjusted payoffs (called “fertilities” by Kroumi and Lessard [2015]) are

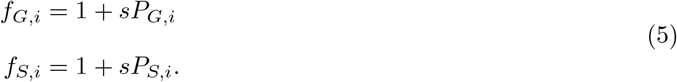

We can obtain fitness by dividing the selection-adjusted payoff against the total selection-adjusted payoff, which is

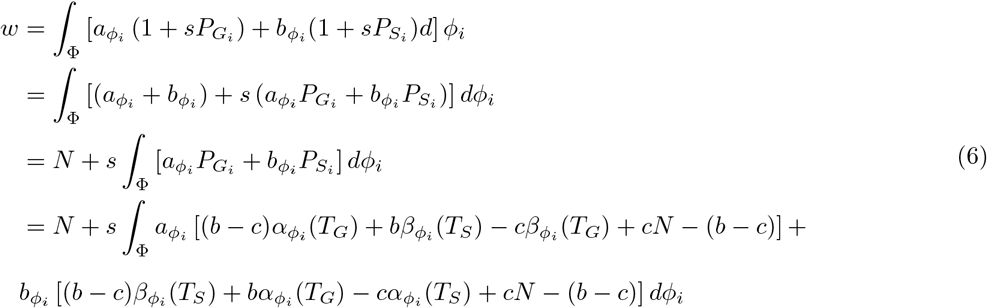

We have as our generous fitness for phenotype *i*

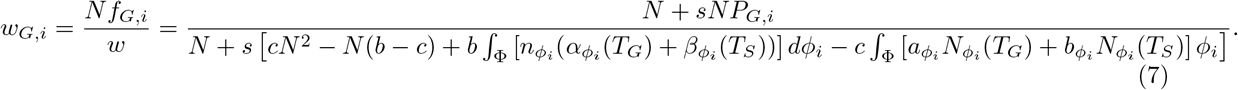

Because we are concerned with the weak selection limit *s* → 0, we perform a first-order Taylor expansion of *w*_*G,i*_ around *s* = 0:

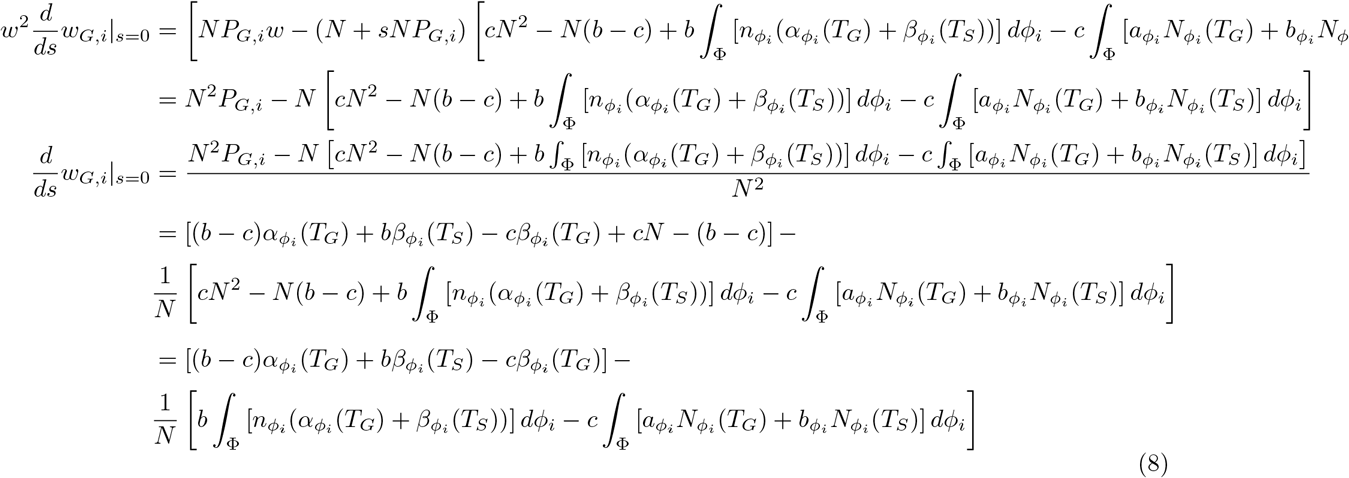

and so

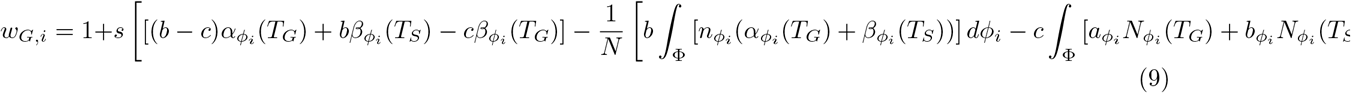

The expected change in frequency due to selection for a particular population state 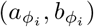 is from eq. 12 in Kroumi and Lessard [2015]:

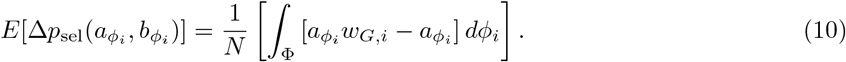

Plugging (9) into (10) yields

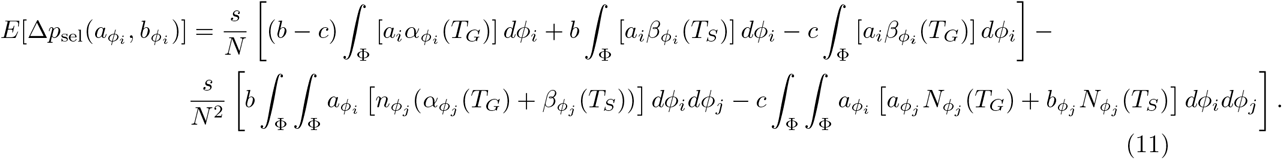

We can take the expectation under 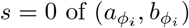 of both sides to obtain

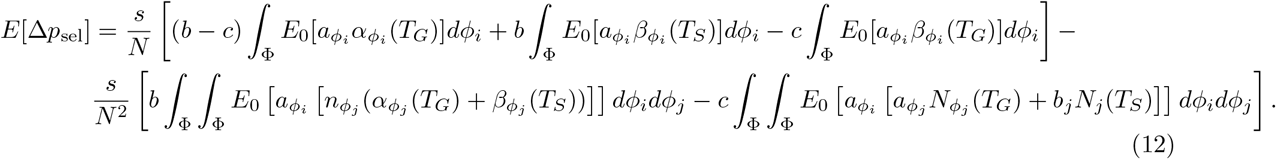

Following Kroumi and Lessard [2015], we define terms according to *I*_*A*_, the indicator function of event *A*. Call P(*i*) the phenotype of individual *i*, and S(*i*) the strategy of individual *i*. Then

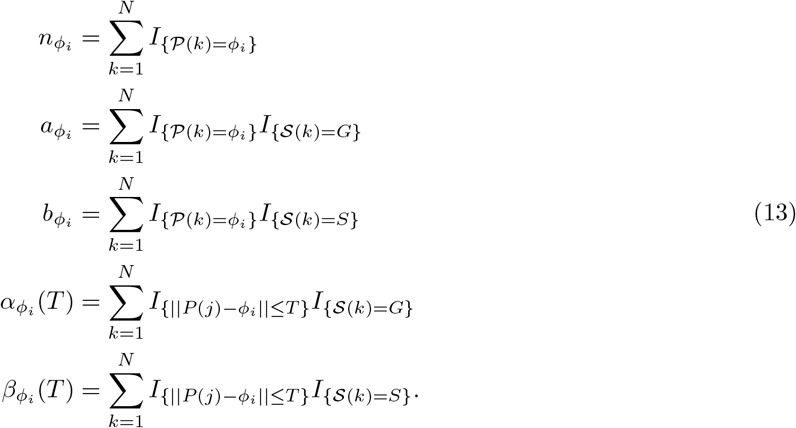

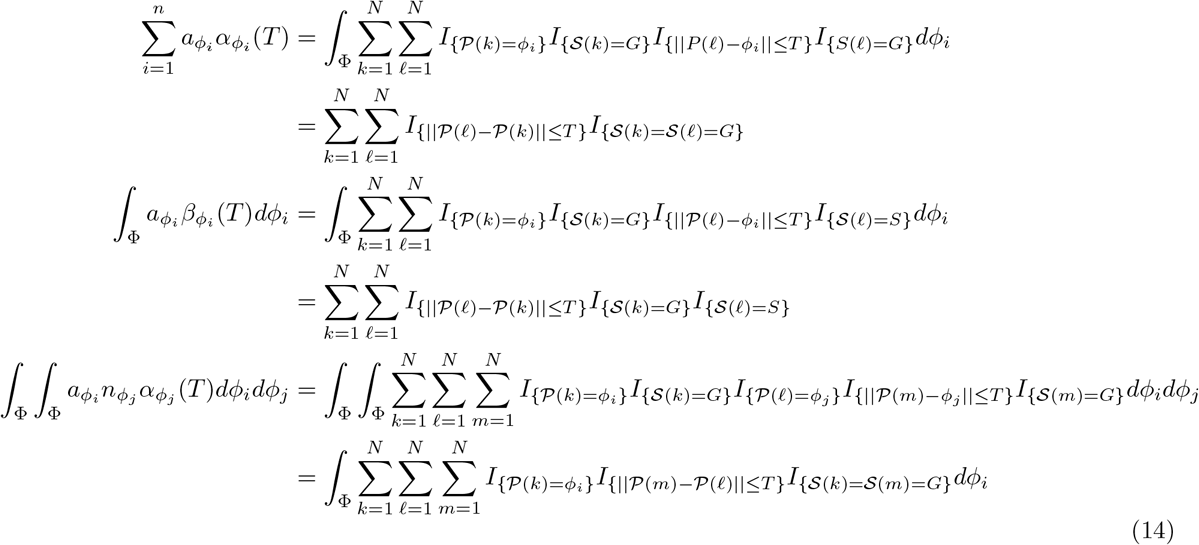

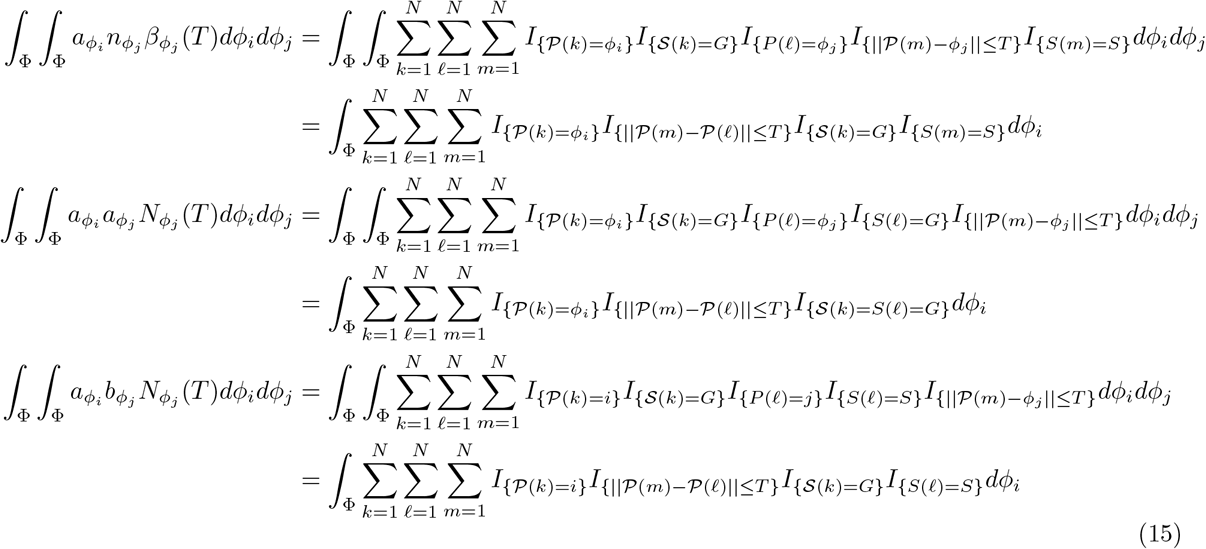

We now define a series of quantities that are analogous to those used by Antal et al. [2009] and Kroumi and Lessard [2015] that describe probabilities relating to three randomly-sampled individuals from the population: *k, f*, and *m*:

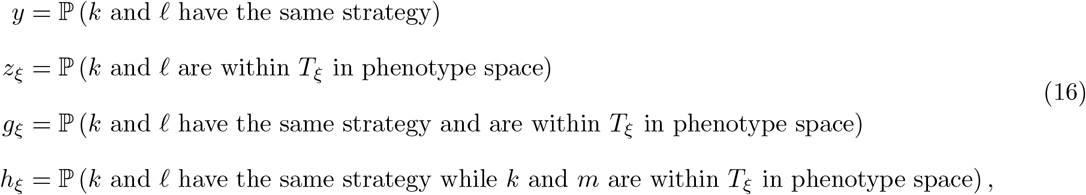

where *ξ* = *G, S* are the strategies.

The two strategies, *G* and *S*, are interchangeable in the neutral model (that is, they should be distributed evenly among the phenotypes), so we have

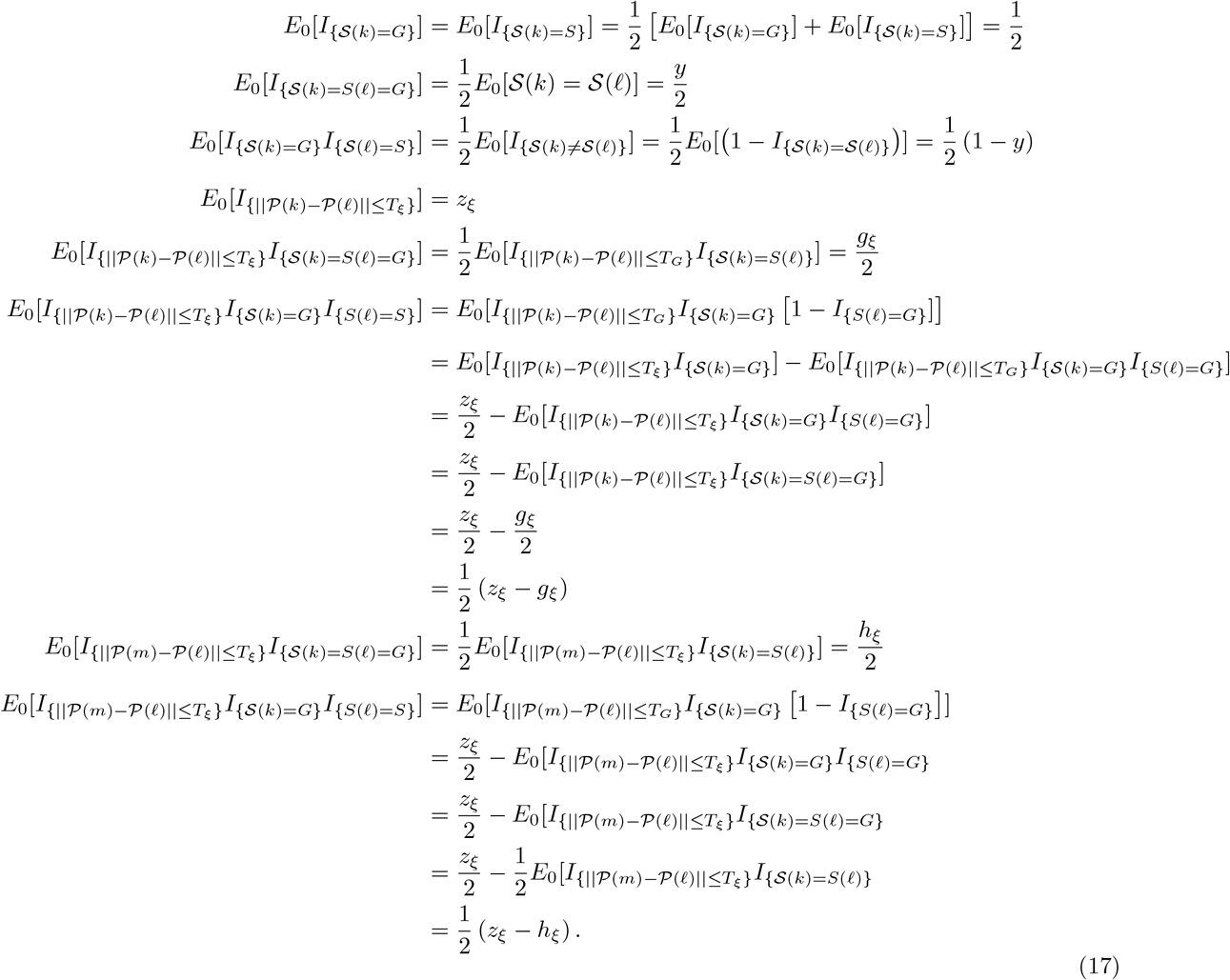

where *ξ* = *G, S*.

Now we can take expectations of the necessary sums:

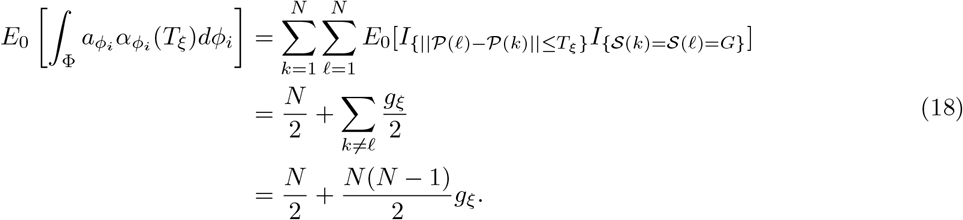

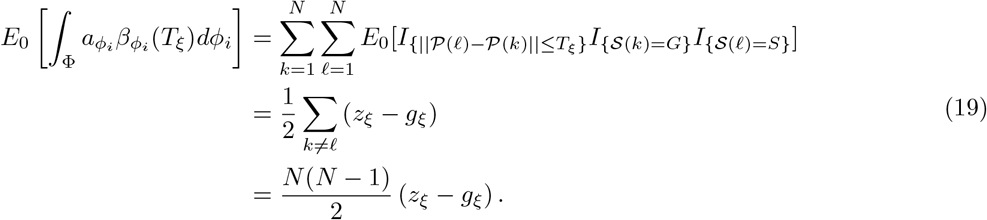

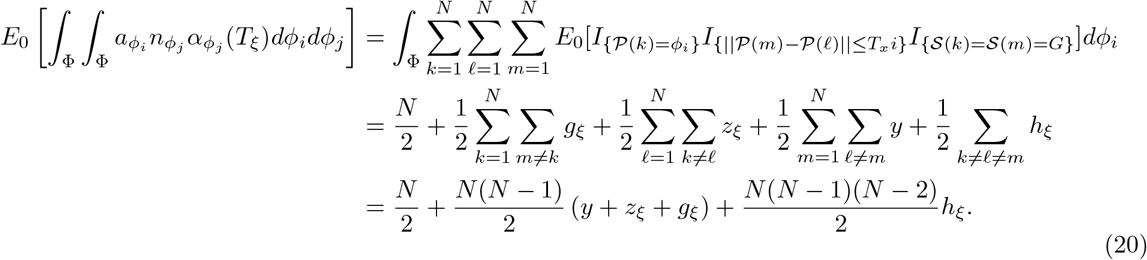

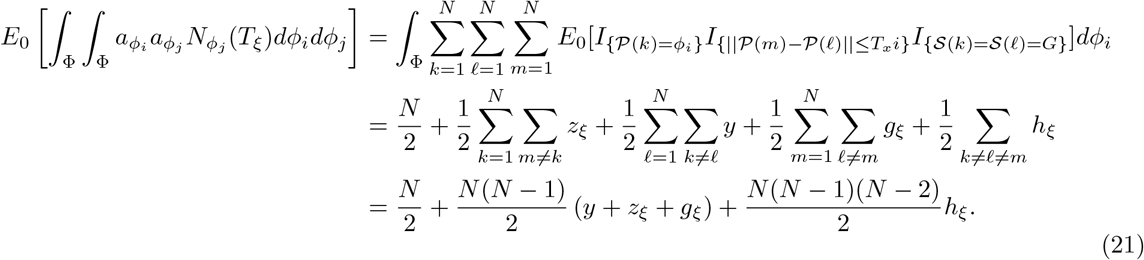

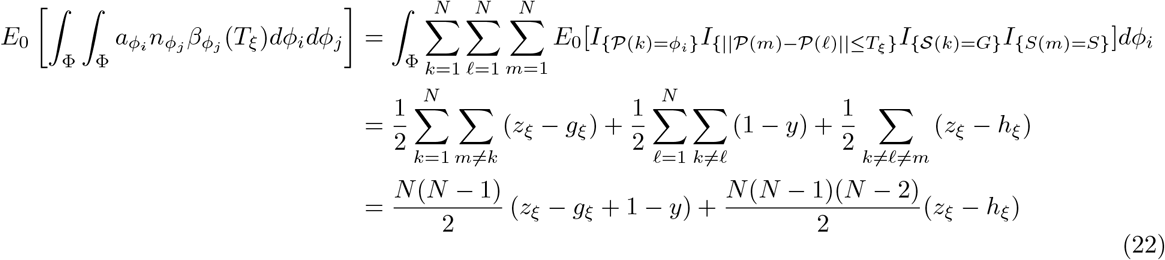

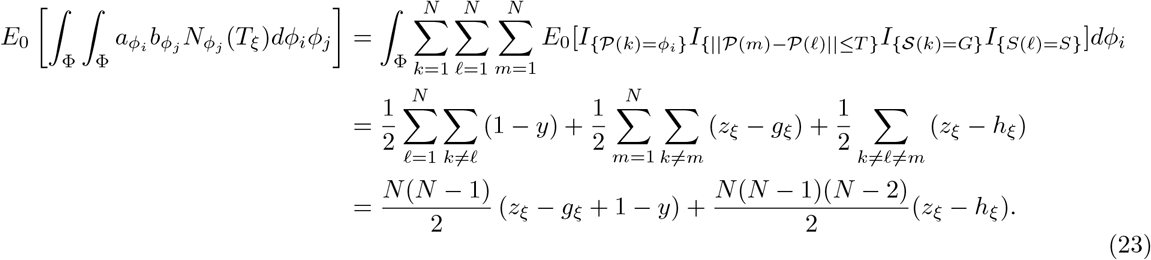

We can now put all of these results into the weak selection expression to obtain the following critical benefit-cost ratio:

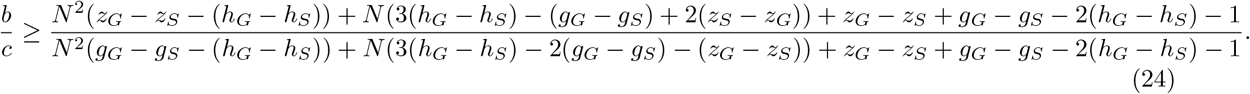

In the limit of *N* → ∞, we have

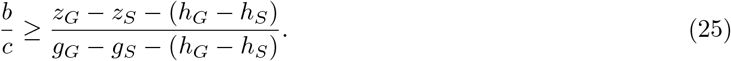

Note that this expression is analogous to eq. 21 from Kroumi and Lessard [2015], which is the same as eq. 49 in Antal et al. [2009].

We now must compute these component terms. We use the same ancestral process as Antal et al. [2009], Kroumi and Lessard [2015], which is a Kingman coalescent in the limit of large population size [Kin] so that we have the same distribution of time to most recent common ancestor of a pair:

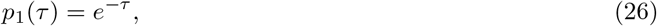

and times to most recent common ancestors for three individuals:

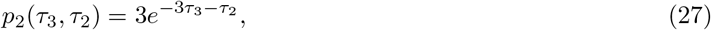

The quantity *μ* = *Nu* is the population strategy mutation rate. If two individuals had a common ancestor *τ* generations ago, then the number of strategy mutations between them is Poisson distributed with rate 2*μτ*. Two individuals have the same strategy if and only if they had an even number of strategy mutations between them. Define *y*(*τ*) to be the probability that two individuals have the same strategy with a most recent common ancestor *τ* generations ago:

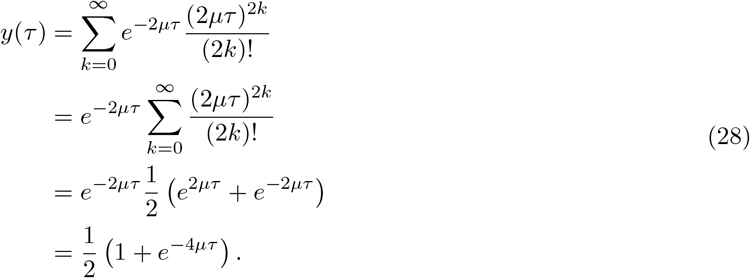

And so we have:

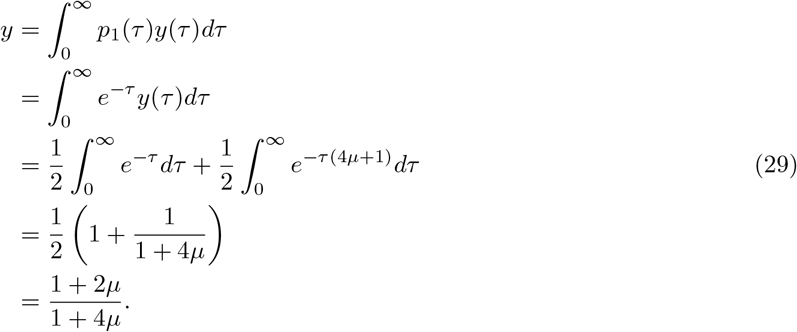

Our strategy mutation model is equivalent to the Kroumi and Lessard [2015] model with their probability of strategy mutation *u*_1_ = 2*u*. Thus, (28) and (29) correspond to eq. 60 and eq. 62 in Kroumi and Lessard [2015], respectively. Note that Antal et al. [2009] use a different definition of *μ*_1_ = 2*Nu*_1_ instead of our 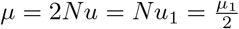, and so (28) and (29) correspond to their eq. 13 and eq. 15, respectively.

To obtain *g* and *h*, we need to consider distances in phenotype space. If two individuals have a most recent common ancestor at time *τ* in coalescent units, then they have had *Nτ* phenotype mutation steps between them. *PM* is the standard deviation of the phenotype mutation size, and we generally set it to 1. Consider the vector that is the difference between the phenotypes of the two individuals. The elements of this vector are differences in individual dimensions of the phenotype. Each such difference is drawn from a normal distribution with mean 0 and variance 2*Nτ* (*PM*)^2^ (the sum of 2 *N* (0, *Nτ* (*PM*)^2^) distributions, which are themselves sums of *Nτ N* (0, (*PM*)^2^) distributions). The magnitude of this difference, the Euclidean distance between these individuals, is therefore *χ*-distributed with dimension *D*, scaled by 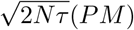: Call 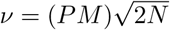 for simplicity’s sake. Then we have:

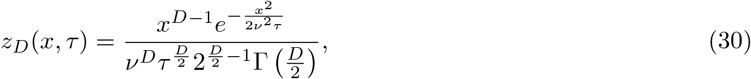

where Γ(*s*) is the Gamma function

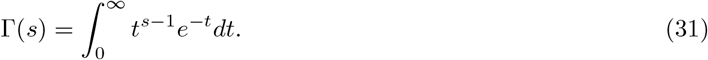

The average pairwise distance for a particular time step *t* (still measuresd in generations) is

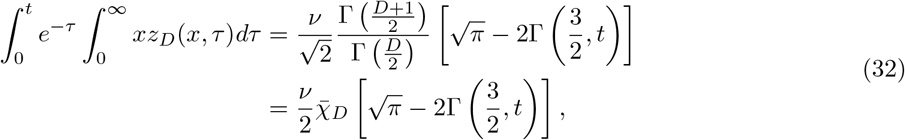

where Γ(*s, k*) is the upper incomplete Gamma function

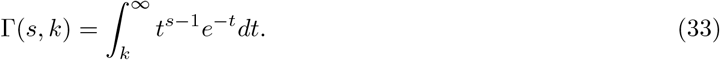

Now we can get the full distribution by integrating over the TMRCAs, which are exponential when time is scaled in coalescent units *τ* :

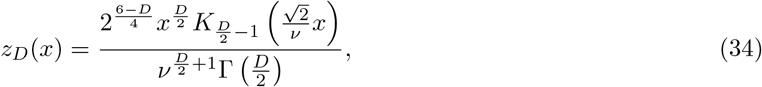

where *D* ≥ 2 and *K*_*α*_(*x*) is the modified Bessel function of the second kind:

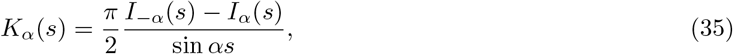

where *I*_*α*_(*s*) is the modified Bessel function of the first kind:

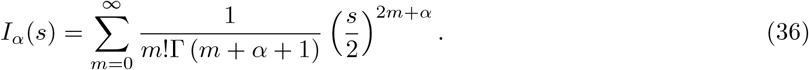

Note that the modified Bessel function of the first kind appears in a similar manner in Kroumi and Lessard [2015].

For *D* = 1, we have instead

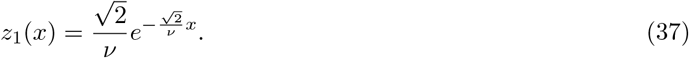

The CDF of (30) over *x* from 0 to *T* is:

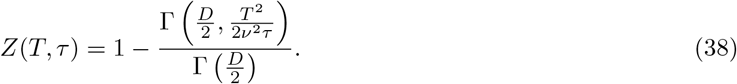

It is useful for ease of notation to define the general quantity:

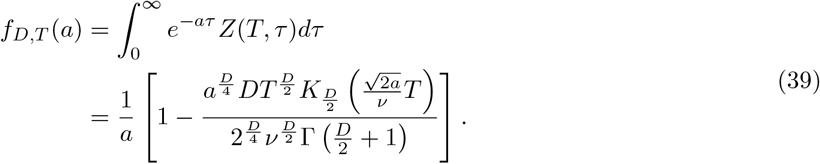

Note that this function is not, in general, a CDF. Also, this function applies for *D* ≥ 2. For *D* = 1, we have

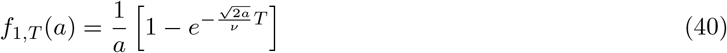

For *a* = 1, we have the CDF of (30) over *x* from 0 to *T* and all *τ* :

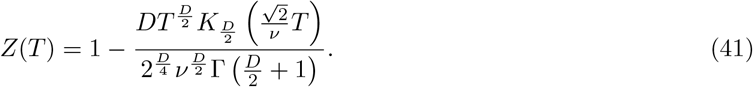

Note that we will in general use *f*_*G*_ and *f*_*S*_ to refer to 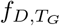 and 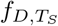, respectively.

The probability of two individuals being within some phenotype distance *T*_*G*_ or *T*_*S*_ is therefore

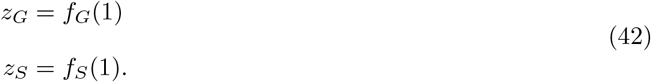

The probability of two individuals being within phenotype distance *T*_*G*_ or *T*_*S*_ and have the same strategy is:

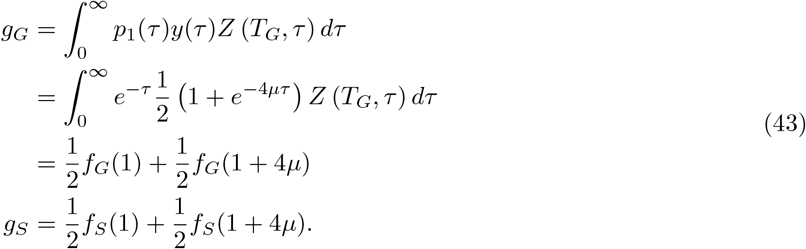

The three-way probability requires the consideration of three different genealogical shapes between the three individuals. Individuals *k* and 𝓁 have the same strategy while *m* and 𝓁 are within *T*_*G*_ of each other. First, let’s say that *m* and 𝓁 coalesce first:

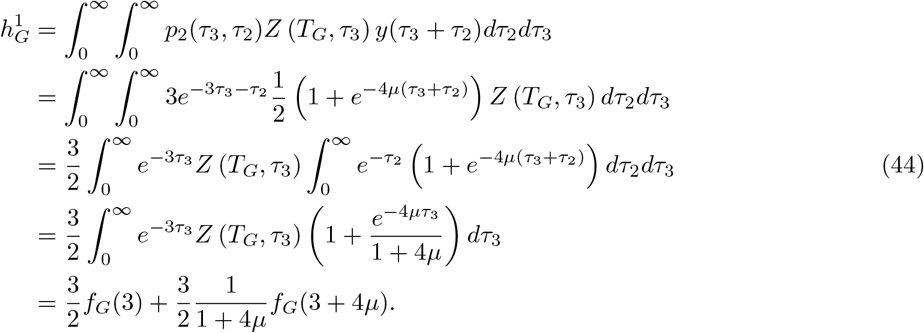

Next, *k* and 𝓁 coalesce first:

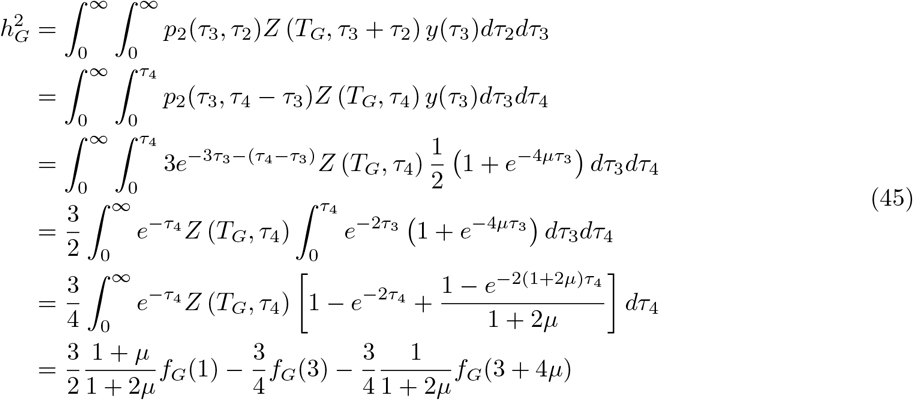

Finally, *k* and *m* coalesce first:

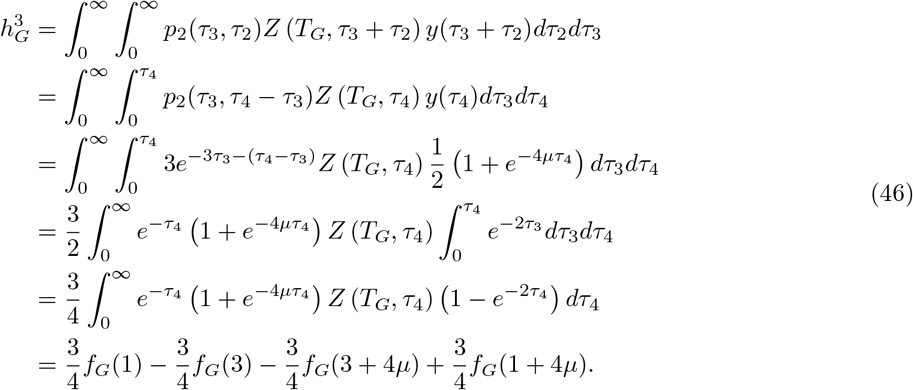

Adding these results together gives us

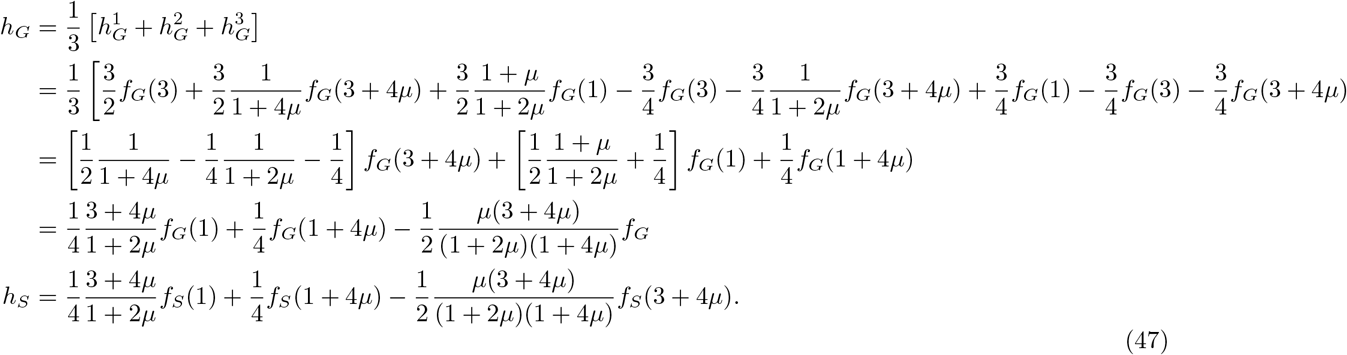

Note that (47) is analogous to eq. 68 in Kroumi and Lessard [2015]. Also note that as *T* → 0, *Z* (*T, τ*) → 0, and so for *T*_*S*_ = 0, *g*_*S*_ = *z*_*S*_ = *h*_*S*_ = 0, and we recover the form of eq. 49 in Antal et al. [2009] and eq. 21 in Kroumi and Lessard [2015]. The actual values of these quantities are different in our model as compared to theirs, but the forms of the equations are the same.

### The neutral phenotype distribution

The mean pairwise phenotypic distance under neutrality can be obtained from (34):

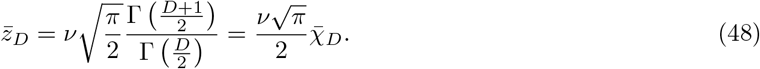

To translate this back into an approximate *χ* distribution, we note that the pairwise distances can be interpreted as distributed as *χ* scaled by 2*σ*^2^, where *σ*_0_ is the standard deviation per dimension. So the standard deviation per dimension must therefore be

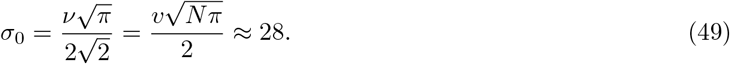

which is close to what we observe and is the value of the horizontal line in Figures 4 and 5.

